# Distinct origin and fate for fetal hematopoietic progenitors

**DOI:** 10.1101/2025.01.08.631951

**Authors:** F. Soares-da-Silva, G. Nogueira, Marie-Pierre Mailhe, L. Freyer, A. Perkins, S. Hatano, Y. Yoshikai, P. Pereira, A. Bandeira, R. Elsaid, E. Gomez-Perdiguero, A. Cumano

## Abstract

It was proposed that two sequential sources of intraembryonic multipotent progenitors ensure blood cell production from late gestation into adulthood, with only the latter producing self-renewing hematopoietic stem cells (HSC). How these two populations differ and how they impact the establishment of the postnatal immune system, remains poorly understood. Using complementary lineage tracing models, we showed that the first emerging embryonic multipotent progenitors (eMPP) are responsible for late gestation hematopoiesis. They are distinct from HSC that do not significantly contribute to embryonic mature blood cells. eMPP are the predominant source of embryonic lymphocytes and lymphoid tissue inducer cells, some of which persist for life. Between E12.5 and E16.5 eMPP rapidly differentiate, whereas HSC expand 20-fold. Altogether, these results support the notion that eMPP establish the embryonic adaptive immune system and shape the lymphoid organs where later adaptive immune responses occur, while HSC expand to sustain blood cell production throughout life.

## Introduction

The hematopoietic system develops from independent, though overlapping waves of progenitors, initially occurring in the yolk sac (YS) vasculature and later in the intra-embryonic dorsal aorta, vitellin and omphalo-mesenteric arteries (Soares-da-Silva et al., 2023; Cumano et al., 2019; Yokomizo and Suda, 2024). Starting at embryonic day (E) 7.5, YS-derived cells initially comprise primitive erythrocytes and later (E8.5-9.5) erythro-myeloid progenitors (EMP) (Palis et al., 1999; Bertrand et al., 2005). EMP generate tissue-resident macrophages that persist throughout life, red blood cells that circulate until birth (Soares-da-Silva et al., 2021) and megakaryocytes, some of which differentiated bypassing the megakaryocyte-erythrocyte progenitor (MEP) stage (Ginhoux et al., 2010; Gomez Perdiguero et al., 2015; Iturri et al., 2021). Hematopoietic stem and progenitor cells (HSPC), comprising multipotent progenitors (MPP) with lymphoid potential, emerge in the major embryonic blood vessels between E9.5 and E11.5 and give rise to the adult hematopoietic stem cell (HSC) compartment that is responsible for life-long blood cell production (Medvinsky and Dzierzak, 1996; Cumano et al., 1996; de Bruijn et al., 2000). Both YS EMP and intra-embryonic HSPC arise through an endothelial to hematopoietic (ETH) transition process (Boisset et al., 2010; Bertrand et al., 2010; Kissa and Herbomel, 2010). Several lines of research suggested that HSPC are diverse in their potential to become adult-type HSC. CXCR4 expression by hemogenic endothelial cells marks progenitors of HSC, whereas CXCR4^-^ cells generate HSPC that do not contribute to adult hematopoiesis (Dignum et al., 2021). Before detection of HSC, Flt3 expression marks multipotent cells, which contribute to fetal hematopoiesis, show a lymphoid differentiation bias, and generate embryonic restricted lymphoid cells (Beaudin et al., 2016). In line with the above studies, lineage tracing experiments in mice showed that fetal hematopoietic cells are independent from HSC and can persist up to 1 year of age (Yokomizo et al., 2022; Kobayashi et al., 2023; Patel et al., 2022).

Whereas all blood lineages are produced throughout life, some specific subsets have been reported to be of exclusive embryonic origin, for example invariant γδ T cells (Havran and Allison, 1990; Heyborne et al., 1993), lymphoid tissue inducer (Lti) cells (Mebius et al., 2001) and B1 cells (Herzenberg, 2000). The first wave of thymic seeding progenitors (TSP), which starts at E12.5, is unique in its capacity to give rise to invariant γδ T and Lti cells, whereas the second wave, first detected at E16.5 in the thymus, only generates conventional αβ and γδ T cells (Ramond et al., 2014; Elsaid et al., 2021). The first TSP are not derived from YS EMP or from IL-7Ra expressing YS progenitors, as suggested (Böiers et al., 2013), but they are labeled together with HSPC in the Csf1r^MER-iCRE-MER^ mouse model pulsed at E10.5 (Elsaid et al., 2021).

To understand the developmental origin of embryonic and adult hematopoietic lineages, we analyzed fetal hematopoiesis in two different inducible lineage tracing models. Induction of Cdh5^CreERT2^ Rosa26^YFP^ mice marks all hemogenic endothelium (Sörensen et al., 2009) and the timing of induction distinguishes different progenitors that emerge at distinct time windows. In contrast, induction of Mds1^CreERT2^ Rosa^26YFP^ mice primarily marks adult-type HSC (Zhang et al., 2021). The analyses of these two models pulsed at multiple time points covering the period from E7.5 to E11.5 allowed the discrimination between the three hematopoietic generations, namely YS-EMP, eMPP, and adult-type HSC. Whereas microglia is of exclusive YS-EMP origin, we show here that invariant γδ T cells, and consequently the first wave of TSP, were of exclusive eMPP origin. In contrast, the second wave of TSP and all other hematopoietic lineages in the adult were derived from adult-type HSC. Although most B1 cells (Herzenberg, 2000) located in the peritoneal cavity of adult mice derived from HSC, a significant contribution from eMPP was observed. Only eMPP contribute to fetal hematopoiesis, whereas by 4 weeks of age, the HSC progeny dominates the hematopoietic compartment. eMPP and HSC co-exist in the FL and cannot be distinguished by currently used surface markers because both contribute to the, phenotypically defined, Lin^-^Sca-1^+^KIT^+^ (LSK) and HSC compartments. However, from E12.5 to E16.5, eMPP rapidly differentiate whereas a reduced number of adult-type HSC expand approximately 20-fold. This work, therefore, allows following the fate of the late waves of hematopoietic cell generation.

## Results

### 1. Detection of transient eMPP before E10.5

Previous work showed that a single injection of OH-tamoxifen (OH-TAM) in Cdh5^CreERT2^ Rosa26^YFP^ mice at E7.5 preferentially marks tissue-resident macrophages and embryonic erythrocytes whereas induction at E10.5 predominantly labels adult-type HSC (Gentek et al., 2018a). However, induction at any of these two days failed to label a significant fraction of thymic invariant γδ T cells (Gentek et al., 2018b), indicating that they derive from hemogenic endothelial cells different from those that give rise to YS EMP or adult-type HSC. This result suggested a transient generation of hematopoietic progenitors that give rise to embryonically restricted lymphoid cells. To identify these progenitors, we injected pregnant Cdh5^CreERT2^ Rosa26^YFP^ females with OH-TAM at E7.5, E8.5, E9.5 and E10.5 and analyzed the embryos at different embryonic days and as adults (Fig 1A). At E16.5, microglia were labeled at high levels after OH-TAM pulse at E7.5 or E8.5 but were no longer labeled by OH-TAM pulses at E9.5 or E10.5 (Fig 1B). Kinetics of YFP-expressing LSK cells in E12.5, E14.5, and E16.5 FL or adult (4-9 weeks post-birth) bone marrow (BM) revealed that LSK in mice pulsed at E7.5, E8.5 or E10.5 had very different fates (Fig 1C, FigS1A). In embryos pulsed with OH-TAM at E7.5, around 10% of the LSK at E14.5 and E16.5, and virtually none by 4 weeks of age, were found labeled, indicating that OH-TAM at E7.5 labels a transient population of LSK (Fig 1C). By contrast, in embryos pulsed at E10.5, the frequency of YFP^+^ LSK had a 2-fold increase between E14.5 and E16.5 and was maintained in the adult. Therefore, OH-TAM at E10.5 labels LSK that contribute to adult hematopoiesis. OH-TAM pulse at E8.5 labeled a much higher frequency of LSK than when pulsed at E7.5 or E10.5 (Fig.S2A). This frequency does not increase between E14.5 and E16.5 and significantly decreases between E16.5 and 4 weeks post-birth (Fig. 1C). The different kinetics of embryos pulsed at E8.5 and E10.5 indicate that OH-TAM at E8.5 labels a heterogeneous population of LSK, some of which contribute to adult hematopoiesis and others that are transient and no longer detected after 4 weeks of post-natal life. This is further confirmed by the fact that the frequency of YFP-expressing LSK in E16.5 FL of Cdh5^CreERT2^ Rosa26^YFP^ mice pulsed at E8.5 and E9.5 is higher than in mice pulsed at E7.5 and E10.5 (FigS2A).

**Figure 1.**
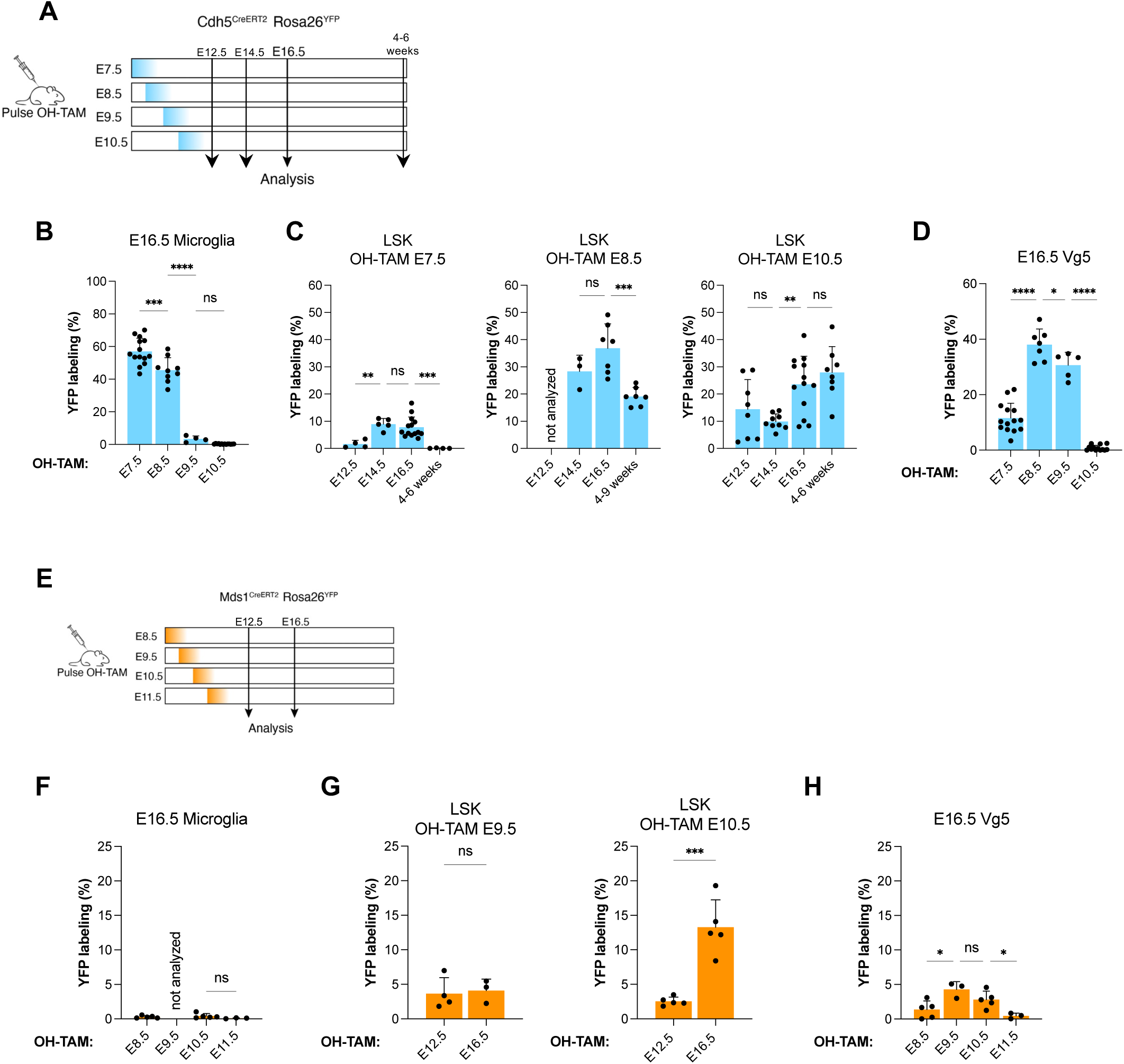
Lineage tracing of LSK have time-dependent dynamics. **A.** Experimental strategy to analyze Cdh5^CreERT2^ Rosa26^YFP^ mice. **B**. Frequency of YFP^+^ cells in E16.5 microglia, from embryos administered OH-TAM at the indicated stages of gestation. **C**. Frequency of YFP^+^ cells in LSK, in FL, at different stages of gestation and at 4-9 weeks of post-natal life of Cdh5^CreERT2^ Rosa26^YFP^ embryos treated with OH-TAM at E7.5 (left panel), E8.5 (middle panel) and E10.5 (right panel). **D**. Frequency of YFP^+^ cells in E16.5 thymic Vγ5 Vδ1 γδ T cells in Cdh5^CreERT2^ Rosa26^YFP^ embryos treated with OH-TAM at the indicated gestational stages. **E**. Experimental strategy to analyze Mds1^CreERT2^ Rosa26^YFP^ mice. **F**. Frequency of YFP+ cells in E16.5 microglia, from embryos administered OH-TAM at the indicated stages of gestation **G**. Frequency of YFP^+^ cells in LSK, in FL, at E12.5 and 16.5 of Mds1^CreERT2^ Rosa26^YFP^ embryos treated with OH-TAM at E9.5 (left panel) and E10.5 (right panel). **H**. Frequency of YFP^+^ cells in E16.5 thymic Vγ5 Vδ1 γδ T cells in Mds1^CreERT2^ Rosa26^YFP^ embryos treated with OH-TAM at the indicated gestational stages. Statistical significance was assessed using one-way ANOVA followed by Šídák’s correction for multiple comparisons. *, P < 0.05; **, P < 0.01; ***, P < 0.001; ****, P < 0.0001. Data are represented as mean ± SD; each dot represents an embryo.

Vγ5Vδ1 T cells, the signature population derived from the first wave of TSP (Ramond et al., 2014; Elsaid et al., 2021), were analyzed in E16.5 thymi of these embryos. In E7.5-pulsed embryos, on average 13% of Vγ5Vδ1 T cells were labeled, a frequency that increased 2.6-fold in E8.5-pulsed embryos (Fig 1D, FigS2A). The frequency of labeled cells was kept very high in E9.5-pulsed embryos, a timepoint when microglia cells were no longer labeled. This distinct kinetics show that Vγ5Vδ1 T cells do not derive from YS-EMP, but from an intermediate wave of HSC-independent progenitors.

Mds1^CreERT2^ Rosa26^YFP^ mice were pulsed at E8.5, E9.5, E10.5 and E11.5 and analyzed at E12.5 or E16.5 (Fig 1E, Fig S2B). Virtually no microglial cells were labeled irrespective of the day of the OH-TAM pulse, confirming that this mouse model does not target YS hemogenic endothelium (Fig 1F) (Zhang et al., 2021). Analyses of LSK in FL at E12.5 and E16.5 revealed that injection of OH-TAM at E8.5 and E9.5 labels a low frequency of LSK that did not increase between the two time points analyzed (Fig. 1G, Fig. S2B). By contrast, OH-TAM at E10.5 labels a population of LSK that increases around 5-fold, in frequency, between E12.5 and E16.5 (Fig. 1G). This result parallels that obtained with the Cdh5^CreERT2^ Rosa26^YFP^ model, indicating that early-labeled LSK are transient (Fig1G, Fig S2B). E16.5 thymi comprised <5% YFP^+^ Vγ5Vδ1^+^ T cells, irrespective of the time of OH-TAM induction, reinforcing the notion that they do not derive from HSC and also shows that this mouse model labels eMPP with a lower frequency than in the Cdh5^CreERT2^Rosa26^YFP^ model. This result is strengthened by the observation that OH-TAM at E10.5 labels more LSK than at E9.5 (Fig.S2B). In conclusion, combining the two mouse models allows tracing YS-EMP, eMPP and adult-type HSC.

### 2. HSC do not contribute to embryonic hematopoiesis

Next, we studied the frequency of YFP-labeled cells in different populations of differentiated or progenitor cells in E16.5 FL and fetal thymi of Cdh5^CreERT2^ Rosa26^YFP^ females injected with OH-TAM at E7.5, E8.5, E9.5 or E10.5 are shown in Fig. 2A. Due to the inherent variability in the frequencies of labeled cells across different pregnant females in independent experiments, frequencies were indexed to those of YFP-labeled LSK in each experiment, hereafter denoted as normalized YFP expression. All mature cells analyzed, such as B lineage cells (CD19^+^), granulocytes (Gr-1^+^) and embryonic restricted Vγ5 or Vγ6 (Hatano et al., 2019) T cells in the thymus, followed a similar pattern of YFP expression (Fig.2A). Injection of OH-TAM at E7.5 resulted in a normalized YFP expression higher than 1, indicating that all these cells could have an HSC-independent, eMPP origin. OH-TAM at E8.5 and 9.5 showed a normalized expression close to 1, whereas OH-TAM at E10.5 showed YFP ratios closer to 0, indicating that differentiated fetal hematopoietic cells did not originate from HSC but rather from eMPP or YS-EMP (Fig.2B). Similar results were obtained in the Mds1^CreERT2^ model (Fig.2D-E).

**Figure 2.**
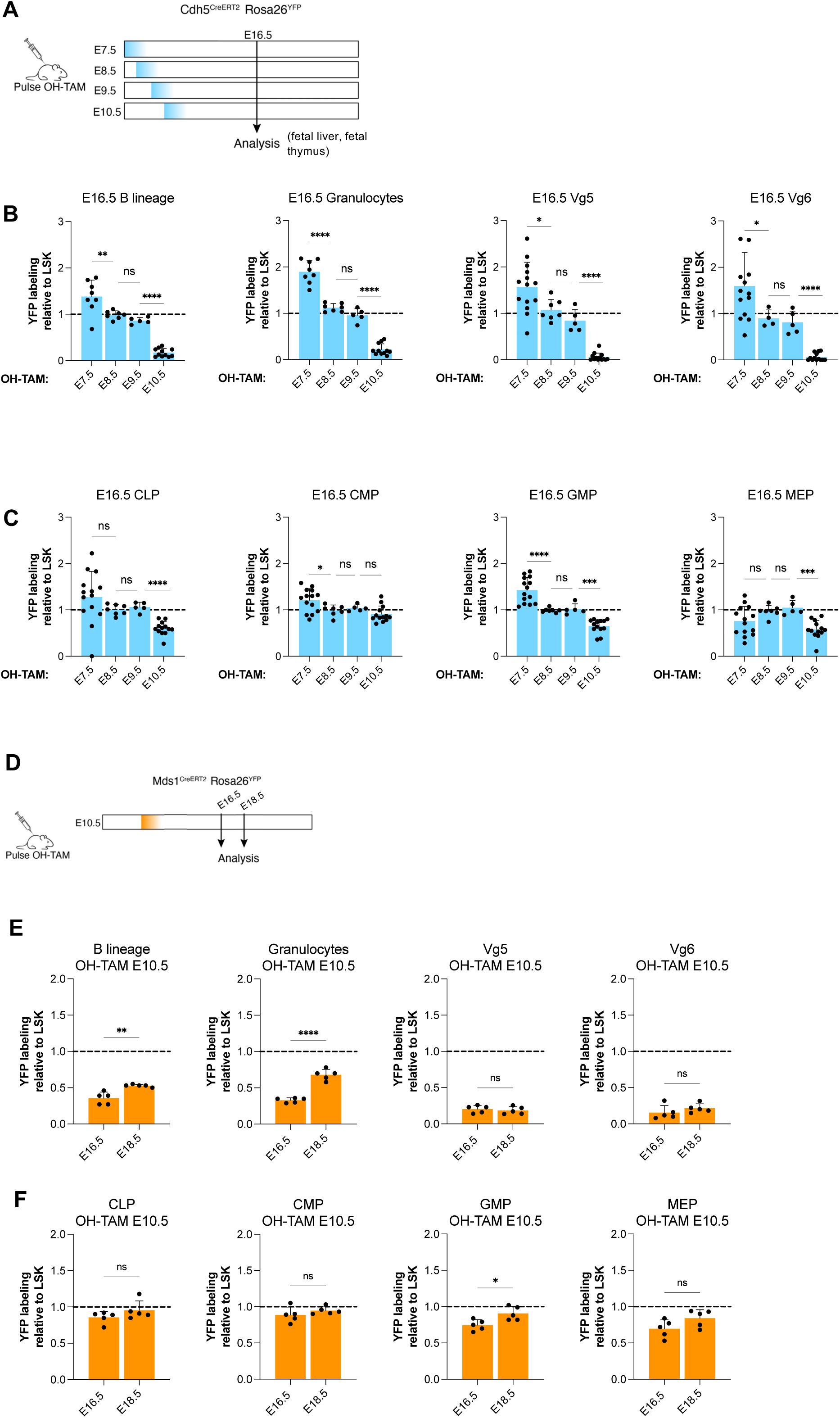
Adult-type HSC do not contribute to the embryonic differentiated compartment. **A.** Experimental design to analyze differentiated and progenitor cells in E16.5 embryos. **B**. Frequency of YFP^+^ cells normalized to the frequency of YFP^+^ LSK, in individual Cdh5^CreERT2^ Rosa26^YFP^ embryos, pulsed with OH-TAM, at the indicated gestational days, and analyzed in the designated differentiated populations (B lineage and granulocytes in FL, Vγ5 and Vγ6 γδ T cells in the thymus). **C**. Frequency of YFP^+^ cells normalized to the frequency of YFP^+^ LSK, in individual Cdh5^CreERT2^ Rosa26^YFP^ embryos, pulsed with OH-TAM at the indicated gestational days and analyzed in the designated populations, in FL. **D**. Experimental design to analyze differentiated and progenitor cells in E16.5 and E18.5 Mds1^CreERT2^ Rosa26^YFP^ embryos. **E**. Frequency of YFP^+^ cells normalized to the frequency of YFP^+^ LSK, in individual Mds1^CreERT2^ Rosa26^YFP^ embryos, pulsed with OH-TAM, at E10.5, and analyzed at E16.5 and E18.5 (B lineage and granulocytes in FL, Vγ5 and Vγ6 γδ T cells in the thymus). **F**. Frequency of YFP^+^ cells normalized to the frequency of YFP^+^ LSK, in individual Mds1^CreERT2^ Rosa26^YFP^ embryos, pulsed with OH-TAM at E10.5 and analyzed at E16.5 and E18.5, in FL. Statistical significance was assessed using one-way ANOVA followed by Šídák’s correction for multiple comparisons. *, P < 0.05; **, P < 0.01; ***, P < 0.001; ****, P < 0.0001. Data are represented as mean ± SD; each dot represents an embryo.

Unlike differentiated cells, the precursor compartment showed normalized YFP values close to 1 for a pulse of OH-TAM at E7.5, 8.5 and 9.5 (Fig 2C). After a pulse of OH-TAM at E10.5, although common myeloid and common lymphoid progenitors (CMP and CLP (Kondo et al., 2001)) were derived from HSC, granulocyte-monocyte progenitors (GMP) and MEP, showed a significant contribution of eMPP, at this stage (E16.5). These results are compatible with the hierarchical progression of hematopoietic progenitors from CMP to GMP and MEP (Akashi et al., 2000) and show that eMPP can contribute to the progenitor compartment, including the LSK and HSC pools in the FL (Fig. S2C). Considering that myeloid cells in FL can derive from YS-EMP, eMPP or HSC (Iturri et al., 2021) we analyzed E10.5-pulsed Mds1^CreERT2^ Rosa26^YFP^ embryos at E16.5 and at E18.5 (Fig. 2D). Consistent with the above results, normalized YFP expression showed that B lineage cells and granulocytes expressed values below 0.5, at E16.5. These frequencies increased at E18.5, indicating that differentiated FL cells, initially of eMPP origin, were gradually replaced by progenies of adult-type HSC (Fig. 2E). As expected, YFP expression in Vγ5 and Vγ6 γδ T cells in the thymus remained constantly low. In the progenitor compartment, YFP expression in CLP and CMP was close to 1, indicating their adult-type HSC origin. GMP and MEP showed normalized YFP expression below 1 at E16.5, which increased at E18.5 indicating that the erythro-myeloid progenitor compartments, which initially comprised the progeny of YS-EMP and eMPP, was dominated by adult-type HSC progeny by the end of gestation (Fig. 2F).

At E16.5, the thymus is mostly composed of thymocytes derived from the first wave of TSP, with only DN1 cells originated from the second wave TSP (Ramond et al., 2014). Cadh5^CreERT2^ Rosa26^YFP^ and Mds1^CreERT2^ Rosa26^YFP^ pulsed with OH-TAM at E10.5, showed a significantly higher YFP ratio in DN1 than in DN3 thymocytes, invariant Vγ5 or Vγ6 T cells, or in Lti cells (Fig S3, left and central panels). These results indicate that, whereas most thymocytes at this stage (E16.5) derive from eMPP, DN1 from the second wave of TSP are HSC-derived. OH-TAM pulse at E8.5, by contrast, exhibits normalized YFP values close to 1 (Fig S3, right panel), compatible with the heterogeneous, bimodal origin of progenitors labeled at this stage.

### 3. Progenies of eMPP do not contribute to hematopoiesis in the adult

To analyze the contribution of eMPP to adult hematopoiesis, 4-9 weeks-old mice were analyzed for the distribution of YFP-labeled cells in the BM, spleen, skin, lung, inguinal lymph nodes (iLN) and peritoneal cavity of Cadh5^CreERT2^ Rosa26^YFP^ mice pulsed with OH-TAM at E8.5 and E10.5 (Fig 3A). Brain microglia were used as an internal control (Fig. 3B). The frequency of YFP-expressing LSK in the BM was significantly increased in E10.5-pulsed mice compared to E8.5-pulsed animals, indicating a higher production of adult-type HSC at E10.5 (Fig.3C). The normalized YFP expression in animals pulsed at E8.5 and E10.5 was close to 1 for splenic T cells, splenic B cells and peritoneal B2 B cells, suggesting their HSC origin at this stage of post-natal life (Fig. 3D-F). By contrast, Vγ5 T cells in the epidermis and Vγ6 T cells in the iLN and lungs showed a ratio close to 0, confirming that they originated from eMPP and not from HSC (Fig.3G). B1a and B1b B cells in the peritoneal cavity showed a YFP ratio lower than 1, indicating that a fraction of them originated from eMPP during fetal life and persist at that stage.

**Figure 3.**
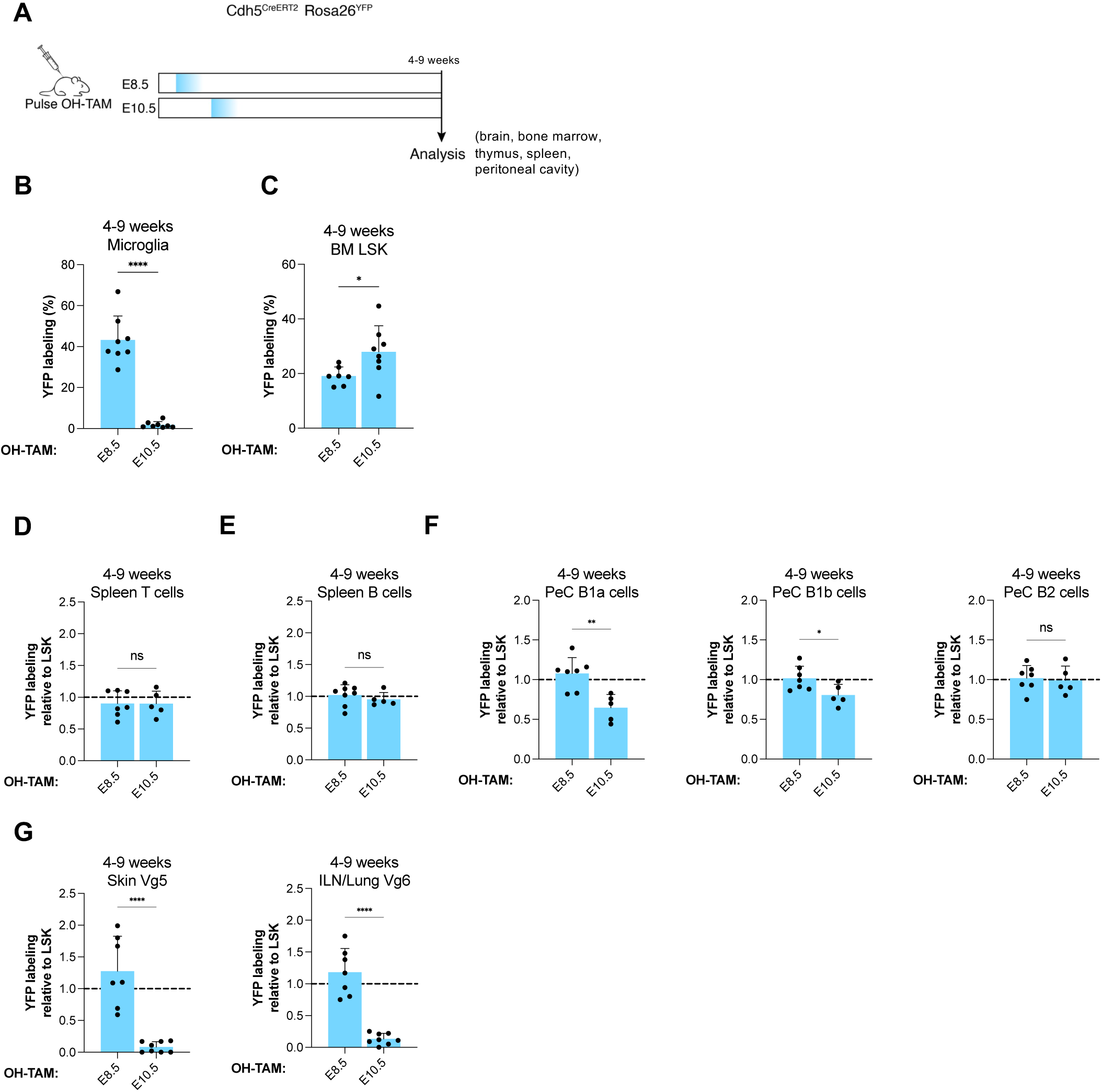
The contribution of E-MPP to adult hematopoiesis is restricted to invariant γδ T cells and to B1 cells, in the peritoneal cavity. **A.** Experimental design to analyze differentiated and progenitor cells in Cdh5^CreERT2^ Rosa26^YFP^ mice at 4-9 weeks post-birth. **B**. Frequency of YFP^+^ microglia in the brain of 4-9 week-old mice, injected with OH-TAM at the indicated gestational days. **C**. Frequency of YFP^+^ cells in BM LSK of 4-9 week-old mice, injected with OH-TAM at the indicated gestational days. **D**. Frequency of YFP^+^ cells, normalized to the frequency of YFP^+^ cells in BM LSK, in splenic T cells of 4-9 week-old mice. **E**. Frequency of YFP^+^ cells, normalized to the frequency of YFP^+^ cells in BM LSK, in splenic B cells of 4-9 week-old mice. **F.** Frequency of YFP^+^ cells, normalized to the frequency of YFP^+^ cells in BM LSK, in the peritoneal cavity of 4-9 week-old mice, B1a B cells (left panel), B1b B cells (middle panel) and B2 B cells (right panel). **G.** Frequency of YFP^+^ cells, normalized to the frequency of BM LSK YFP^+^ cells in epidermal Vγ5^+^ γδ T cells of 4-9 week old mice (left panel), or Vγ6 γδ T cells in the inguinal lymph nodes or lungs of 4-9 week old mice (right panel). Statistical significance was assessed using one-way ANOVA followed by Šídák’s correction for multiple comparisons. *, P < 0.05; **, P < 0.01; ***, P < 0.001; ****, P < 0.0001. Data are represented as mean ± SD; each dot represents an embryo.

These results demonstrate a unique contribution of transient hematopoietic progenitors to the production of embryonically restricted lymphoid populations and B1a and B1b cells.

### 4. HSC expand around 20-fold in the fetal liver from E12.5 to E16.5

A recent study (Ganuza et al., 2022) suggested that HSC, exclusively defined by phenotype, undergo minimal expansion in the FL. However, the fact that HSC and eMPP fall in the same LSK gate formally precludes that conclusion. As shown in Fig.S2C, both subsets show a similar distribution regarding the two markers, indicating that transient eMPP integrate these two LSK compartments. Therefore, phenotypically defined progenitors in the FL are heterogeneous in origin and fate, with a fraction of cells that derive from eMPP and rapidly differentiate and a fraction that generates the adult HSC compartment.

Because the Mds1^CreERT2^ model preferentially labels adult-type HSC while labeling eMPP with low frequency, we analyzed progenitor expansion in this model. We first estimated the numbers of phenotypically defined LSK, and HSC in the FL (Fig 4A) of Cdh5^CreERT2^ Rosa26^YFP^, Mds1^CreERT2^ Rosa26^YFP^ and their littermate controls, between E12.5 and E16.5.

**Figure 4.**
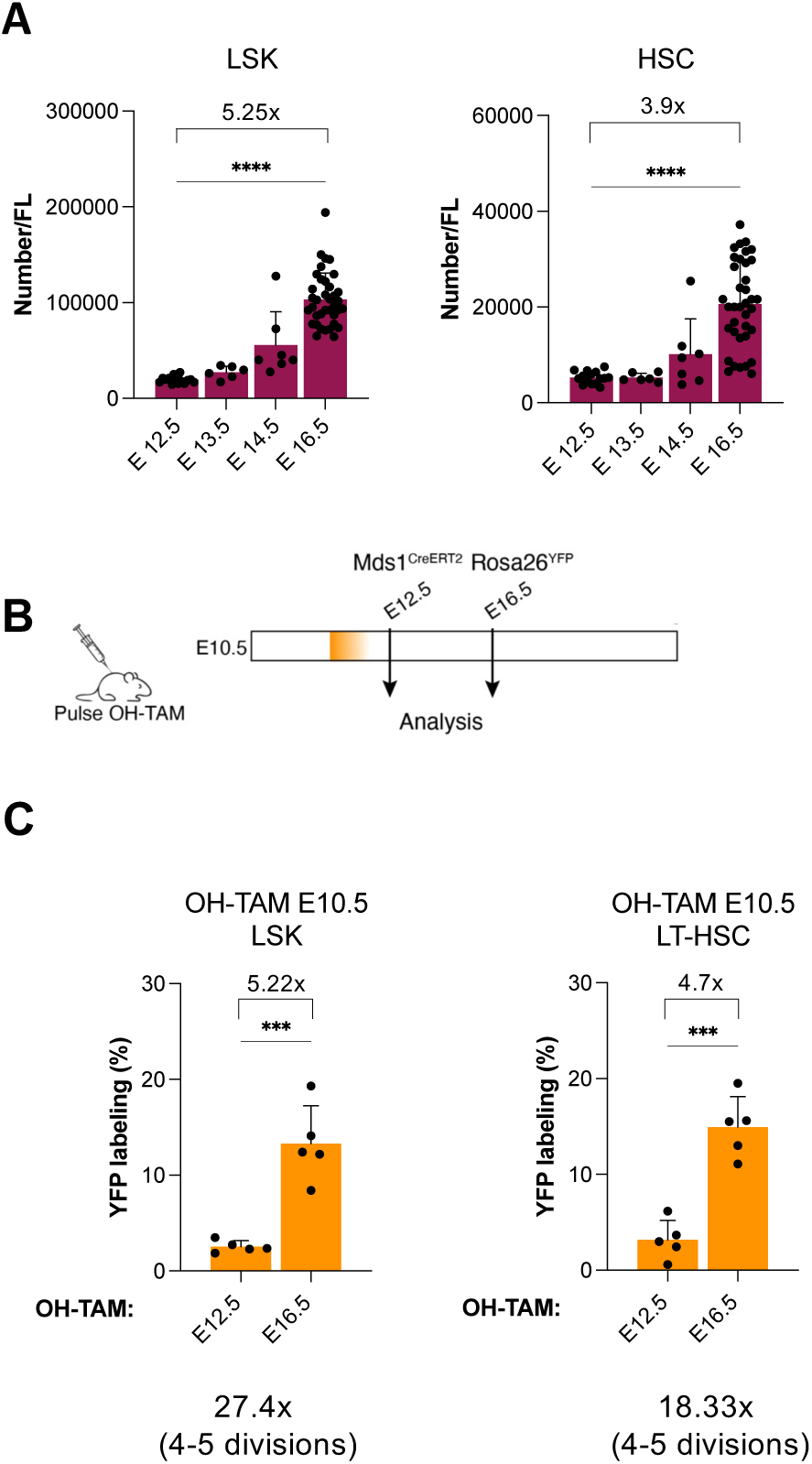
LSK and adult-type HSC undergo significant expansion, in FL. **A**. Numbers of phenotypically defined LSK and HSC (LSK CD48^-^CD150^+^) calculated at different gestational stages in FL. **B**. Experimental design to analyze differentiated and progenitor cells in Mds1^CreERT2^ Rosa26^YFP^ mice E12.5 and E16.5. **C.** Frequencies of YFP^+^ at E12.5 and E16.5 in embryos pulsed with OH-TAM at E10.5. Statistical significance was assessed using one-way ANOVA followed by Šídák’s correction for multiple comparisons. *, P < 0.05; **, P < 0.01; ***, P < 0.001; ****, P < 0.0001. Data are represented as mean ± SD; each dot represents an embryo.

We found that LSK and HSC (LSK CD48^-^CD150^+^) numbers increased around 5,3-fold and 3,9-fold, respectively from E12.5 to E16.5 (Fig 4A). Analyses of E10.5-pulsed Mds1^CreERT2^ Rosa26^YFP^ mice showed a 4,7-fold expansion of HSC from E12.5 to E16.5 (Fig. 4B,C) and a similar expansion (5,2-fold) of total LSK. Similar analyses in mice pulsed at E9.5 showed no increase in the fractions of labeled total LSK and HSC suggesting that eMPP do not expand during this time window (Fig S4A). The net fold increase in HSC between E12.5 and E16.5 can be calculated as the product of the increase in the number of HSC by the increase in the frequency of YFP^+^ cells. When calculated for total LSK and HSC, the net fold increase was 27- and 18-fold, respectively (Fig 4C), what corresponds to 4-5 rounds of division within 4 days. To directly analyze cell proliferation in vivo Cdh5^CreERT2^ Rosa26^YFP^ mice pulsed at E9.5 (to mark cells of eMPP mixed origin) or at E10.5 (to preferentially mark HSC) were injected with a single dose of EdU at E13.5 and analyzed 90 minutes later for EdU^+^ cells in YFP^+^ FL LSK that were CD48^+^ or CD48^-^ (Fig S4B). About 50% and 30% of the cells were EdU^+^ in the CD48^+^ and CD48^-^ compartments, respectively (Fig. S4B).

The results were similar in the two OH-TAM induction time-points. Since eMPP do not increase in number during this period, these results support the notion that they rapidly differentiate. By contrast, the extensive proliferation and increase in numbers of HSC demonstrates their expansion in the FL.

## Discussion

The continuous generation of HSPC generation requires tight control of the timing of induction of lineage tracing and barcoding experiments. Our data support this notion by showing that induction of Cre in Cdh5^CreERT2^ Rosa26^YFP^ mice at E8.5 and E9.5 labels a population that contains YS-EMP, eMPP and adult-type HSC in unknown ratios highlighting the largely overlapping nature of the EHT process. Our observations also indicate that, throughout fetal life, the FL harbors two distinct but phenotypically indistinguishable populations of HSPC. One restricted to embryonic life consisted of eMPP that contributes to fetal but not adult hematopoiesis. The other population expanded, without immediate differentiation, to form the adult pool of HSC. This work is in line with other studies (Inlay et al., 2014; Beaudin and Forsberg, 2016; Dignum et al., 2021; Yokomizo et al., 2022), with the exception that we did not detect a significant contribution of eMPP to conventional hematopoiesis in post-natal life (Patel et al., 2022). At early gestational stages (E12.5), phenotypically defined HSC are mostly eMPP that will soon differentiate, whereas a small number are adult-type HSC that expand and will dominate the phenotypically defined HSC compartment by late gestation. Studies that consider fetal progenitors as a homogeneous population should be reinterpreted with this new perspective. By using two complementary lineage tracing models, our study allows the distinction between the three hematopoietic generations (YS-EMP, eMPP and HSC), to follow their fate and track a population enriched in adult-type HSC. We show that the latter undergo around 20-fold expansion in the FL from E12.5 to E16.5, corresponding to 4 to 5 rounds of division. This value assumes no differentiation or cell death and is, therefore, a minimal estimate.

The unique embryonic nature of certain lymphoid populations may result from the fact that they originate from progenitors different from those of adult lymphocytes. These embryonic lymphoid cells, also described in humans (Tan et al., 2021), persist in low numbers throughout life and function to maintain tissue integrity (Guo et al., 2018) and promote wound healing (Havran and Jameson, 2010). These findings suggest that embryonic lymphoid and myeloid cells may have substantially different development and function from their adult counterparts and might be at the origin of disorders mainly affecting infants (Horton et al., 2013). Indeed, childhood leukemias were shown to arise during fetal life and are infrequent after puberty (Greaves, 2018). We propose that the eMPP traced in this study represent the target population for the mutations/translocations that initiate acute lymphoblastic leukemia in infants.

## Supplemental Figures

**Figure S1.**
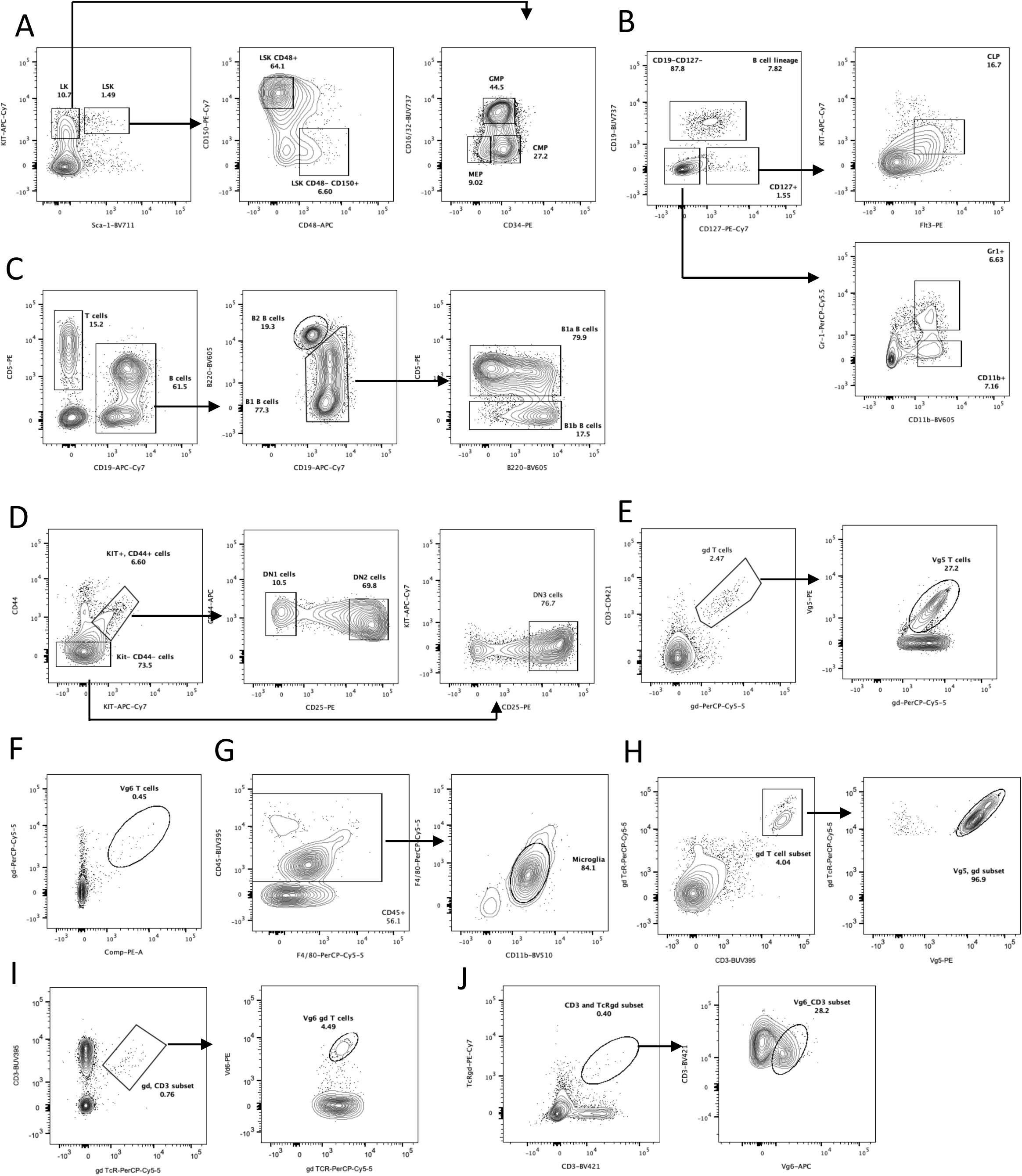
Gating strategies to identify different cell populations in the embryo and in adult mice. FL cells, peritoneal cells or thymocytes were gated for singletons and live cells **A**. Cells were stained with biotinylated antibodies for lineage (TER119, CD71, CD19, CD3, Gr-1, NK-1.1, CD11C), CD16/32, CD34, CD45, KIT, Sca-1, CD48, CD150 and streptavidin to define lineage^+^ cells. This staining defining CD45^+^ LSK, HSC (LSK CD48^-^ and CD150^+^), CMP (LK, CD16/32^-^CD34^+^), GMP (LK, CD16/32^+^CD34^+^), MEP (LK, CD16/32^-^CD34^-^). **B**. TER119^-^CD71^-^CD45^+^CD19^+^ defines B lineage cells; CD19^-^ CD127^+^KIT^interm^, Flt3^+^ define CLP; TER119^-^ CD71^-^CD45^+^CD19^-^CD127^-^CD11b^-^Gr-1^+^ define granulocytes. **C**. CD11b^-^, CD19, IgM and B220+ cells define B cells. B220^high^CD19^+^ defines B2 B cells, B220^interm/low^ CD19^+^ defines B1 B cells.CD5 defines B1a (CD5^+^) and B1b (CD5^-^) B cells. **D**. Thymocytes were stained with biotinylated CD3, CD4 and CD8 antibodies, CD44, CD25, KIT and streptavidin. CD3^-^CD4^-^CD8^-^CD44^+^ KIT^+^ defined early thymic progenitors (ETP), CD25^-^ ETP are DN1 and CD25^+^ are DN2. **E**. Thymocytes were stained with γδ, CD3 Vγ5 antibodies to define Vγ5 expressing γδ T cells. **F**. CD3, γδ and anti-Vγ6 staining was followed by anti-rat IgG to define Vγ6 γδ T cells. **G**. Brain single cell suspensions were stained with anti-CD45 and the positive fraction was analyzed for the expression of F4/80 and CD11b defining microglia. **H**. Single cell suspensions of ear epidermis were analyzed inr CD3/γδTcR positive cells for Vγ5 expression. **I**. Inguinal lymph node suspensions were stained with anti-CD3 and anti-γδ TcR and Vγ6 γδ T cells were identified using the anti-Vγ6 antibody followed by a secondary staining with anti-rat IgG. **J**. Single cell suspensions from lungs were stained with anti-CD3, anti-TcRγδ and anti-Vγ6.

**Figure S2.**
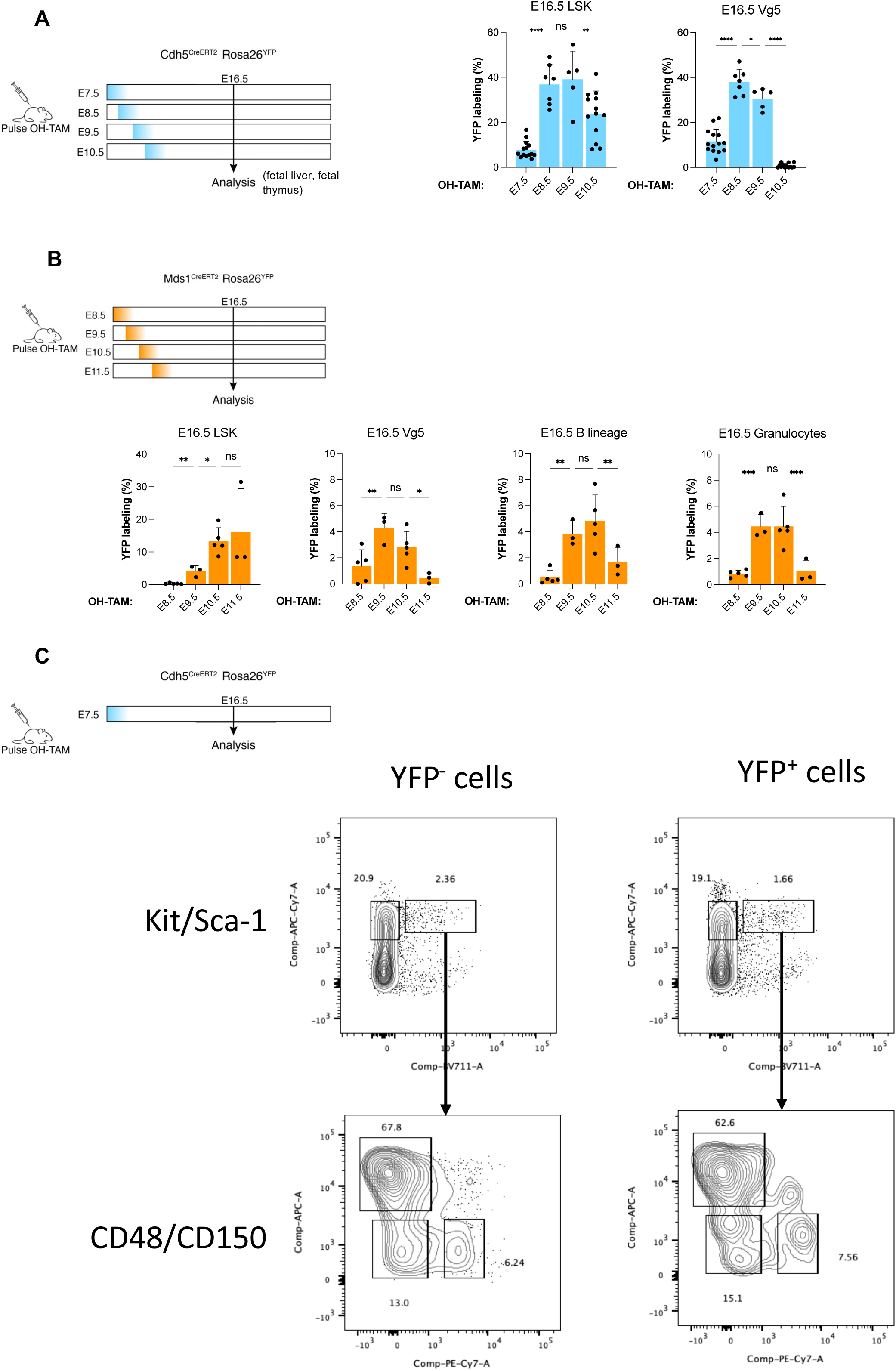
Frequencies of YFP^+^ cells in embryonic cell compartments. **A**. Frequencies of YFP^+^ cells in E16.5 embryos of Cdh5^CreERT2^ Rosa26^YFP^ pulsed with OH-TAM, at the indicates gestational stages, in FL LSK (left panel) and Vγ5^+^ γδ T cells in the thymus (right panel). **B**. Frequencies of YFP^+^ cells in E16.5 embryos of Mds1^CreERT2^ Rosa26^YFP^ embryos pulsed with OH-TAM, at the indicated gestational stages, in FL LSK (left panel), thymic Vγ5^+^ T cells (middle left panel), FL B lineage cells (middle right panel) and FL granulocytes (right panel). **C**. Frequency of LSK, CD48^+^CD150^-^, CD48^-^CD150^+^ and CD48^+^CD150^+^ in YFP^+^ and YFP^-^ E16.5 FL cells of Cdh5^CreERT2^ Rosa26^YFP^ embryos pulsed with OH-TAM at E7.5. Statistical significance was assessed using one-way ANOVA followed by Šídák’s correction for multiple comparisons. *, P < 0.05; **, P < 0.01; ***, P < 0.001; ****, P < 0.0001. Data are represented as mean ± SD; each dot represents an embryo.

**Figure S3.**
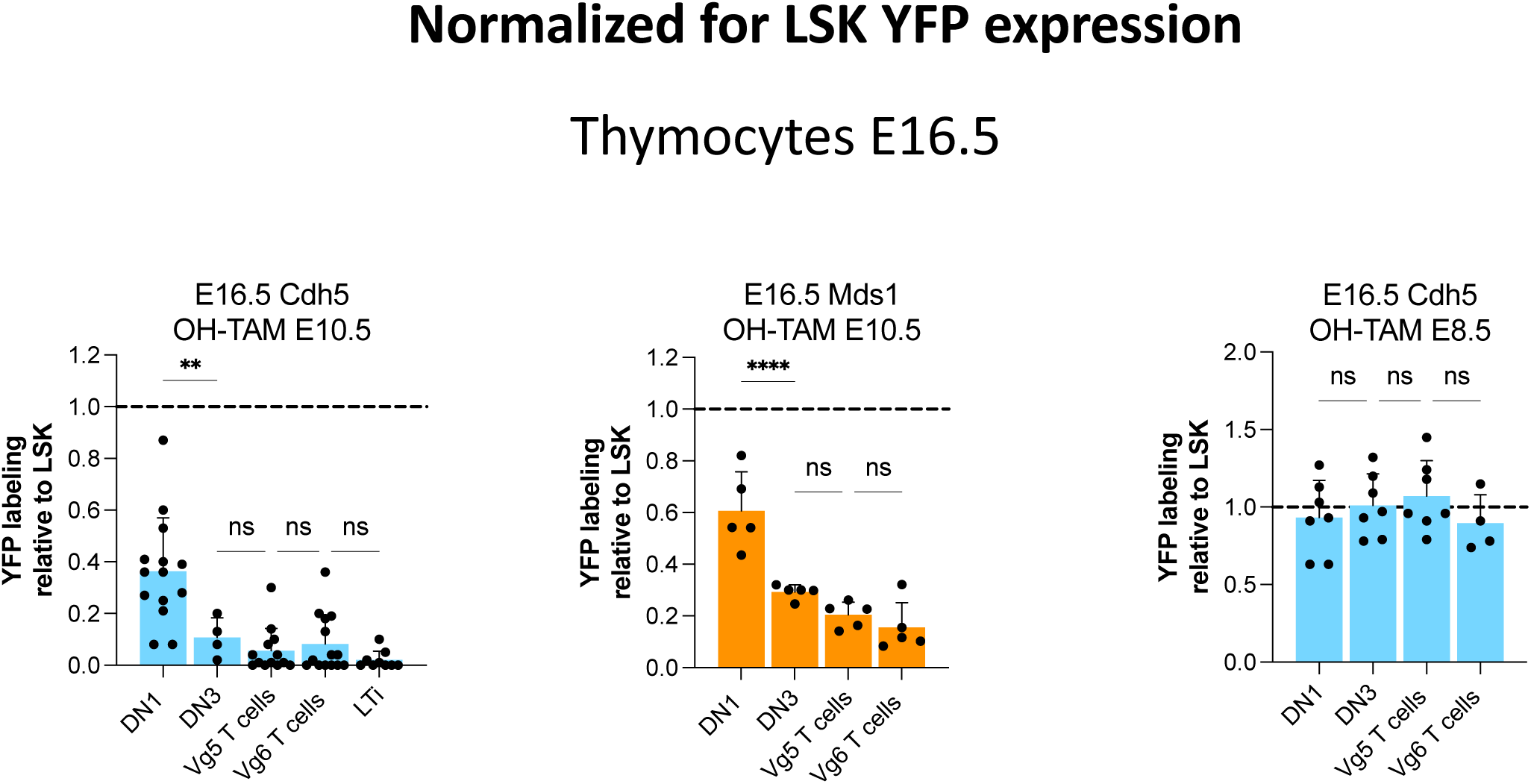
The second wave of TSP originate from adult-type HSC. Frequency of YFP^+^ cells normalized to the frequency of YFP^+^ BM LSK in the designated E16.5 thymocyte subsets of Cdh5^CreERT2^ Rosa26^YFP^ embryos pulsed with OH-TAM at E10.5 (left panel), of Mds1^CreERT2^ Rosa26^YFP^ embryos pulsed with OH-TAM at E10.5 (middle panel) and Cdh5^CreERT2^ Rosa26^YFP^ pulsed with OH-TAM at E8.5 (right panel). Statistical significance was assessed using one-way ANOVA followed by Šídák’s correction for multiple comparisons. *, P < 0.05; **, P < 0.01; ***, P < 0.001; ****, P < 0.0001. Data are represented as mean ± SD; each dot represents an embryo.

**Figure S4.**
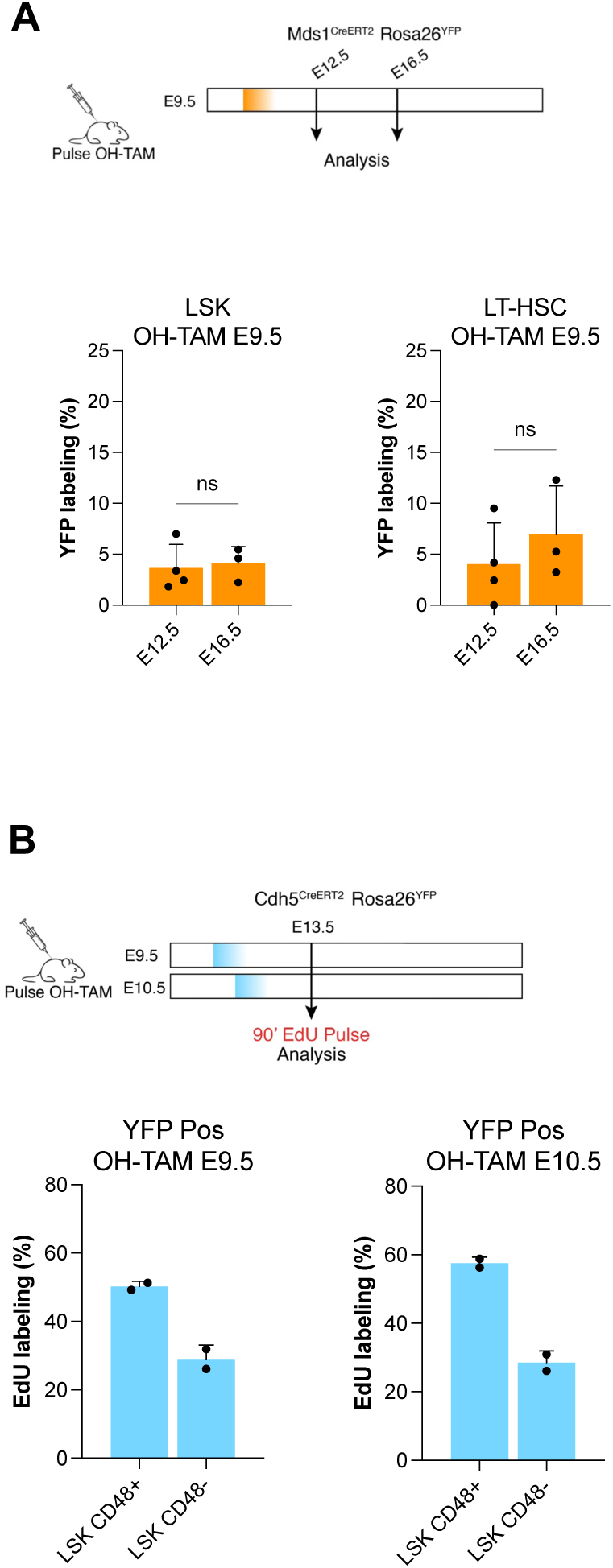
Adult-type HSC preferentially expand in FL. **A**. Frequency of YFP^+^ cells in FL LSK and HSC, at the designated stages, in Mds1^CreERT2^ Rosa26^YFP^ embryos pulsed with OH-TAM at E9.5. **B**. Frequency of EdU^+^ cells in YFP^+^ LSK CD48^+^ or LSK CD48^-^ cells of Cdh5^CreERT2^ Rosa26^YFP^ embryos pulsed with OH-TAM at E9.5 (left panels) or at E10.5 (right panel), in E13.5 FL. Statistical significance was assessed using one-way ANOVA followed by Šídák’s correction for multiple comparisons. *, P < 0.05; **, P < 0.01; ***, P < 0.001; ****, P < 0.0001. Data are represented as mean ± SD; each dot represents an embryo.

## Acknowledgments

We acknowledge the Center for Translational Science (CRT) - Cytometry and Biomarkers Unit of Technology and Service (CB UTechS) at Institut Pasteur for support in conducting this study, namely S. Novault, S. Megharba, S. Schmutz; and the staff of the animal facility of the Institut Pasteur for mouse care.

## AUTHOR CONTRIBUTIONS

F.S.S., E.G.P., R.E., and A.C. designed the experiments; F.S.S., E.G.P., R.E., G.N., and A.C. co-wrote the manuscript; F.S.S., R.E., L.F., M.M. and G.N. performed experiments; A.P., S.H., and Y.Y., provided reagents; P.P, and A.B., contributed to the discussion, designed and performed experiments; and all authors contributed to the manuscript.

## Star Methods

### Mice

C57BL/6J mice were purchased from Envigo. Cdh5^CreERT2^ mice (Sörensen et al., 2009) were provided by Elisa Gomez-Perdiguero (Institut Pasteur). Mds1^CreERT2^ mice (Zhang et al., 2021) were provided by Archibald Perkins (Rochester University). Cdh5^CreERT2^ and Mds1^CreERT2^ mice were crossed to Rosa26^YFP^ reporter mice, kindly provided by Gerard Eberl (Institut Pasteur). 6 to 8-week-old mice or timed pregnant females were used. Timed pregnancies were generated after overnight mating, and the following morning females with vaginal plugs were considered to be at E0.5. The day of birth was considered to be P0. Pregnant females were sacrificed by CO_2_ inhalation, and cervical dislocation as a confirmation method. Cdh5^CreERT2^ Rosa26^YFP^ and Mds1^CreERT2^ Rosa26^YFP^ pregnant females were injected intra-peritoneally with a single dose of 37.5 μg/g body weight of OH-Tamoxifen (Sigma-Aldrich, Cat# H7904) and 18.75 mg/g body weight of progesterone (Sigma-Aldrich, Cat# P3972) between E7.5 and E11.5. For analysis of embryos, pregnant females were sacrificed between E12.5-E16.5. Decapitation was performed for embryos older than E15.5. For analysis of mice between 4-9 weeks, c-section was performed at E19.5 and pups were raised with foster mothers. All animal manipulations were performed according to the ethics charter approved by the French Agriculture Ministry and to the European Parliament Directive 2010/63/EU.

### Flow cytometry

#### Cell suspensions

For the isolation of hematopoietic cells, E12.5-E18.5 FL were dissected under a binocular magnifying lens, recovered in HBSS with 1% FCS (Gibco), and passed through a 26-gauge needle of a 1-mL syringe to obtain single-cell suspensions. Before staining, cell suspensions were filtered with a 100 μm cell strainer (BD). Embryonic thymi were processed similarly to FLs.

Femurs and tibias were isolated from adult mice to isolate BM hematopoietic populations, and surrounding tissues were removed. Bones were flushed using a 26-gauge needle of a 1-ml syringe with HBSS with 1% FCS (Gibco). BM clumps were re-suspended by gentling aspirating with a 19-gauge needle, filtered with a 100 μm cell strainer (BD) and stained.

For microglia analysis, brains were collected and digested for 30 min at 37°C with 0.5 ml of digestion solution as in (Iturri et al., 2021).

For adult skin, ears were harvested from adult mice, the dorsal and ventral sheets were separated using forceps and the epidermis was removed by incubation for 30 min at 37°C in 2.4 mg/ml of Dispase II (Invitrogen). The tissue was then disrupted with forceps to get a single-cell suspension and filtered with a 100 μm cell strainer (BD).

For lung collection, mice were exsanguinated and the lung was minced and incubated for 30 min at 37°C with agitation in RPMI containing 2% FCS, then incubated for 1 hour with RPMI medium containing 0.2 mg/ml Collagenase IV (GIBCO) and 0.1 mg/ml DNase I (Roche). Cells were resuspended in a 70% (vol/vol) solution of Percoll topped by a 40% (vol/vol) solution of Percoll. After 20 min of centrifugation at 3000 rpm, cells were collected at the 40%-70% interface.

Peritoneal washouts of adult mice were performed by injecting PBS intra-peritoneally with HBSS with 1% FCS and collecting the resulting peritoneal fluid.

Lymph nodes were processed by mechanical disruption using a 100 μm cell strainer (BD).

#### Surface marker staining and acquisition

Cell suspensions were stained with the antibodies listed in **Table S1** for 20 min at 4°C. Cells were analyzed on a custom BD LSR Fortessa (BD Biosciences) according to the guidelines for the use of FC and cell sorting (Cossarizza et al., 2021). Dead cells were excluded by incorporation of Propidium Iodide (PI). Data were analyzed with FlowJo software (v.10.8.2, BD Biosciences).

### EdU incorporation and cell cycle analysis

Cdh5^CreERT2^ Rosa26^YFP^ mice were injected intra-peritoneally with EdU at E13.5 and sacrificed 90 minutes post-injection. EdU detection was performed following the manufacturer instructions using the Click-iT EdU Pacific blue flow cytometry assay kit (Invitrogen; Cat# C10418).

### Statistical analysis

The results are presented as mean ± standard deviation (SD). Statistical analyses were conducted using GraphPad Prism, with the specific tests employed indicated in the figure legends.

---

**Table S1.** List of antibodies used for Flow Cytometry (FC).

| Protein | Conjugation | Clone | Catalog # | Manufacturer | Species | Used for the detection of |
| --- | --- | --- | --- | --- | --- | --- |
| B220 | BV605 | RA3-6B2 | 553708 | BD Biosciences | Rat | B Lineage Cells |
| B220 | Biotin | RA3-6B2 | 1116020 | SONY | Rat | Lineage depletion |
| CD117 (KIT) | APC-Cy7 | 2B8 | 1129130 | SONY | Rat | Hematopoietic Progenitors |
| CD127 | PE-Cy7 | A7R34 | 1275070 | SONY | Rat | Hematopoietic Progenitors |
| CD11b | BV510 | M1/70 | 562127 | BD Biosciences | Rat | Myeloid Lineage Cells |
| CD11b | BV605 | M1/70 | 101205 | SONY | Rat | Myeloid Lineage Cells |
| CD11b | PB | M1/70 | 1106120 | SONY | Rat | Myeloid Lineage Cells |
| CD11b | Biotin | M1/70 | 1106020 | SONY | Rat | Myeloid Lineage Cells |
| CD11c | Biotin | N418 | 1186520 | SONY | Rat | Lineage Depletion |
| CD150 | PE-Cy7 | TC15-12F12.2 | 1179570 | SONY | Rat | Hematopoietic Progenitors |
| CD16/32 | BUV737 | 2.4G2 | 6127783 | BD Biosciences | Rat | Hematopoietic Progenitors |
| F4/80 | PerCP-Cy5.5 | BM8 | 1215640 | SONY | Rat | Macrophages |
| CD24 | BV510 | M1/69 | 110915 | SONY | Rat | Erythroid Cells |
| CD34 | PE | RAM34 | 551387 | BD Biosciences | Rat | Hematopoietic Progenitors |
| CD3e | Biotin | 145-2C11 | 1101520 | SONY | Rat | Lineage Depletion |
| CD3e | PB | 500A2 | 558214 | BD Biosciences | Rat | T cells |
| CD3e | BUV395 | 145-2C11 | 563565 | BD Biosciences | Rat | T cells |
| CD4 | Biotin | GK1.5 | 1102020 | SONY | Rat | Lineage Depletion |
| CD4 | BV786 | RM4.5 | 563727 | BD Biosciences | Rat | T cells |
| CD45 | BUV395 | 30-F11 | 564279 | BD Biosciences | Rat | Hematopoietic Cells |
| CD48 | APC | HM48-1 | 1117060 | SONY | Rat | Hematopoietic Progenitors |
| CD71 | BV510 | C2 | 563112 | BD Biosciences | Rat | Erythroid Cells |
| CD71 | Biotin | RI7217 | 1169015 | SONY | Rat | Lineage Depletion |
| CD8a | Biotin | 53-6.7 | 1103520 | SONY | Rat | Lineage Depletion |
| CD8a | PE-Cy7 | 53-6.7 | 552877 | BD Biosciences | Rat | T cells |
| Gr-1 | PerCP-CY5.5 | RB6-8C5 | 1142140 | SONY | Rat | Granulocytes |
| Gr-1 | Biotin | RB6-8C5 | 553125 | BD Biosciences | Rat | Lineage Depletion |
| NK1.1 | Biotin | PK136 | 1143520 | SONY | Rat | Lineage Depletion |
| Sca1 | BV711 | D7 | 1140655 | SONY | Rat | Hematopoietic Progenitors |
| Ter119 | Biotin | TER-119 | 1181020 | SONY | Rat | Lineage Depletion |
| CD44 | APC | IM7 | 559250 | BD Biosciences | Rat | Thymocytes |
| CD25 | PE | PC61 | 553866 | BD Biosciences | Rat | Thymocytes |
| TCR$\gamma\delta$ | PERCP-CY5.5 | GL3 | 1190590 | SONY | Rat | T cells |
| Vγ5 | PE | 5D3 | 553230 | BD Biosciences | Rat | T cells |
| Vγ6 | Unlabeled | 1C10-1F7 | - | - | Rat | T cells |
| CD5 | PE | 53-7.3 | 100608 | SONY | Rat | T and B1a B cells |
| CD19 | BUV737 | 1D3 | 612781 | BD Biosciences | Rat | B cells |
| CD19 | APC-Cy7 | 1D3 | 551655 | BD Biosciences | Rat | B cells |
| CD19 | Biotin | 6D5 | 1177520 | SONY | Rat | B cells |
| Flt3 | PE | A2F10.1 | 553842 | BD Biosciences | Rat | Hematopoietic progenitors |
| Streptavidin | BV785 | - | 2626245 | SONY | - |  |
| Streptavidin | PB | - | S11222 | Invitrogen | - |  |
| Anti Mouse-IgG | PE | - | 550589 | BD Biosciences | Goat | Secondary antibody |

## References

Akashi, K., Traver, D., Miyamoto, T., and Weissman, I. L. (2000). A clonogenic common myeloid progenitor that gives rise to all myeloid lineages. Nature 404, 193–197.

Beaudin, A. E., Boyer, S. W., Perez-Cunningham, J., Hernandez, G. E., Derderian, S. C., Jujjavarapu, C., Aaserude, E., MacKenzie, T., and Forsberg, E. C. (2016). A Transient Developmental Hematopoietic Stem Cell Gives Rise to Innate-like B and T Cells. Cell Stem Cell 19, 768–783.

Beaudin, A. E., and Forsberg, E. C. (2016). To B1a or not to B1a: do hematopoietic stem cells contribute to tissue-resident immune cells. Blood 128, 2765–2769.

Bertrand, J. Y., Chi, N. C., Santoso, B., Teng, S., Stainier, D. Y., and Traver, D. (2010). Haematopoietic stem cells derive directly from aortic endothelium during development. Nature 464, 108–111.

Bertrand, J. Y., Jalil, A., Klaine, M., Jung, S., Cumano, A., and Godin, I. (2005). Three pathways to mature macrophages in the early mouse yolk sac. Blood 106, 3004–3011.

Böiers, C., Carrelha, J., Lutteropp, M., Luc, S., Green, J. C., Azzoni, E., Woll, P. S., Mead, A. J., Hultquist, A., Swiers, G., Perdiguero, E. G., Macaulay, I. C., Melchiori, L., Luis, T. C., Kharazi, S., Bouriez-Jones, T., Deng, Q., Pontén, A., Atkinson, D., Jensen, C. T., Sitnicka, E., Geissmann, F., Godin, I., Sandberg, R., de Bruijn, M. F., and Jacobsen, S. E. (2013). Lymphomyeloid contribution of an immune-restricted progenitor emerging prior to definitive hematopoietic stem cells. Cell Stem Cell 13, 535–548.

Boisset, J. C., van Cappellen, W., Andrieu-Soler, C., Galjart, N., Dzierzak, E., and Robin, C. (2010). In vivo imaging of haematopoietic cells emerging from the mouse aortic endothelium. Nature 464, 116–120.

Cossarizza, A., Chang, H. D., Radbruch, A., Abrignani, S., Addo, R., Akdis, M., Andrä, I., Andreata, F., Annunziato, F., Arranz, E., Bacher, P., Bari, S., Barnaba, V., Barros-Martins, J., Baumjohann, D., Beccaria, C. G., Bernardo, D., Boardman, D. A., Borger, J., Böttcher, C., Brockmann, L., Burns, M., Busch, D. H., Cameron, G., Cammarata, I., Cassotta, A., Chang, Y., Chirdo, F. G., Christakou, E., Čičin-Šain, L., Cook, L., Corbett, A. J., Cornelis, R., Cosmi, L., Davey, M. S., De Biasi, S., De Simone, G., Del Zotto, G., Delacher, M., Di Rosa, F., Di Santo, J., Diefenbach, A., Dong, J., Dörner, T., Dress, R. J., Dutertre, C. A., Eckle, S. B. G., Eede, P., Evrard, M., Falk, C. S., Feuerer, M., Fillatreau, S., Fiz-Lopez, A., Follo, M., Foulds, G. A., Fröbel, J., Gagliani, N., Galletti, G., Gangaev, A., Garbi, N., Garrote, J. A., Geginat, J., Gherardin, N. A., Gibellini, L., Ginhoux, F., Godfrey, D. I., Gruarin, P., Haftmann, C., Hansmann, L., Harpur, C. M., Hayday, A. C., Heine, G., Hernández, D. C., Herrmann, M., Hoelsken, O., Huang, Q., Huber, S., Huber, J. E., Huehn, J., Hundemer, M., Hwang, W. Y. K., Iannacone, M., Ivison, S. M., Jäck, H. M., Jani, P. K., Keller, B., Kessler, N., Ketelaars, S., Knop, L., Knopf, J., Koay, H. F., Kobow, K., Kriegsmann, K., Kristyanto, H., Krueger, A., Kuehne, J. F., Kunze-Schumacher, H., Kvistborg, P., Kwok, I., Latorre, D., Lenz, D., Levings, M. K., Lino, A. C., Liotta, F., Long, H. M., Lugli, E., MacDonald, K. N., Maggi, L., Maini, M. K., Mair, F., Manta, C., Manz, R. A., Mashreghi, M. F., Mazzoni, A., McCluskey, J., Mei, H. E., Melchers, F., Melzer, S., Mielenz, D., Monin, L., Moretta, L., Multhoff, G., Muñoz, L. E., Muñoz-Ruiz, M., Muscate, F., Natalini, A., Neumann, K., Ng, L. G., Niedobitek, A., Niemz, J., Almeida, L. N., Notarbartolo, S., Ostendorf, L., Pallett, L. J., Patel, A. A., Percin, G. I., Peruzzi, G., Pinti, M., Pockley, A. G., Pracht, K., Prinz, I., Pujol-Autonell, I., Pulvirenti, N., Quatrini, L., Quinn, K. M., Radbruch, H., Rhys, H., Rodrigo, M. B., Romagnani, C., Saggau, C., Sakaguchi, S., Sallusto, F., Sanderink, L., Sandrock, I., Schauer, C., Scheffold, A., Scherer, H. U., Schiemann, M., Schildberg, F. A., Schober, K., Schoen, J., Schuh, W., Schüler, T., Schulz, A. R., Schulz, S., Schulze, J., Simonetti, S., Singh, J., Sitnik, K. M., Stark, R., Starossom, S., Stehle, C., Szelinski, F., Tan, L., Tarnok, A., Tornack, J., Tree, T. I. M., van Beek, J. J. P., van de Veen, W., van Gisbergen, K., Vasco, C., Verheyden, N. A., von Borstel, A., Ward-Hartstonge, K. A., Warnatz, K., Waskow, C., Wiedemann, A., Wilharm, A., Wing, J., Wirz, O., Wittner, J., Yang, J. H. M., and Yang, J. (2021). Guidelines for the use of flow cytometry and cell sorting in immunological studies (third edition). Eur J Immunol 51, 2708–3145.

Cumano, A., Berthault, C., Ramond, C., Petit, M., Golub, R., Bandeira, A., and Pereira, P. (2019). New Molecular Insights into Immune Cell Development. Annu Rev Immunol 37, 497–519.

Cumano, A., Dieterlen-Lievre, F., and Godin, I. (1996). Lymphoid potential, probed before circulation in mouse, is restricted to caudal intraembryonic splanchnopleura. Cell 86, 907–916.

de Bruijn, M. F., Speck, N. A., Peeters, M. C., and Dzierzak, E. (2000). Definitive hematopoietic stem cells first develop within the major arterial regions of the mouse embryo. EMBO J 19, 2465–2474.

Dignum, T., Varnum-Finney, B., Srivatsan, S. R., Dozono, S., Waltner, O., Heck, A. M., Ishida, T., Nourigat-McKay, C., Jackson, D. L., Rafii, S., Trapnell, C., Bernstein, I. D., and Hadland, B. (2021). Multipotent progenitors and hematopoietic stem cells arise independently from hemogenic endothelium in the mouse embryo. Cell Rep 36, 109675.

Elsaid, R., Meunier, S., Burlen-Defranoux, O., Soares-da-Silva, F., Perchet, T., Iturri, L., Freyer, L., Vieira, P., Pereira, P., Golub, R., Bandeira, A., Perdiguero, E. G., and Cumano, A. (2021). A wave of bipotent T/ILC-restricted progenitors shapes the embryonic thymus microenvironment in a time-dependent manner. Blood 137, 1024–1036.

Ganuza, M., Hall, T., Myers, J., Nevitt, C., Sánchez-Lanzas, R., Chabot, A., Ding, J., Kooienga, E., Caprio, C., Finkelstein, D., Kang, G., Obeng, E., and McKinney-Freeman, S. (2022). Murine foetal liver supports limited detectable expansion of life-long haematopoietic progenitors. Nat Cell Biol 24, 1475–1486.

Gentek, R., Ghigo, C., Hoeffel, G., Bulle, M. J., Msallam, R., Gautier, G., Launay, P., Chen, J., Ginhoux, F., and Bajénoff, M. (2018a). Hemogenic Endothelial Fate Mapping Reveals Dual Developmental Origin of Mast Cells. Immunity 48, 1160–1171.e5.

Gentek, R., Ghigo, C., Hoeffel, G., Jorquera, A., Msallam, R., Wienert, S., Klauschen, F., Ginhoux, F., and Bajénoff, M. (2018b). Epidermal γδ T cells originate from yolk sac hematopoiesis and clonally self-renew in the adult. J Exp Med 215, 2994–3005.

Ginhoux, F., Greter, M., Leboeuf, M., Nandi, S., See, P., Gokhan, S., Mehler, M. F., Conway, S. J., Ng, L. G., Stanley, E. R., Samokhvalov, I. M., and Merad, M. (2010). Fate mapping analysis reveals that adult microglia derive from primitive macrophages. Science 330, 841–845.

Gomez Perdiguero, E., Klapproth, K., Schulz, C., Busch, K., Azzoni, E., Crozet, L., Garner, H., Trouillet, C., de Bruijn, M. F., Geissmann, F., and Rodewald, H. R. (2015). Tissue-resident macrophages originate from yolk-sac-derived erythro-myeloid progenitors. Nature 518, 547–551.

Greaves, M. (2018). A causal mechanism for childhood acute lymphoblastic leukaemia. Nat Rev Cancer 18, 471–484.

Guo, X. J., Dash, P., Crawford, J. C., Allen, E. K., Zamora, A. E., Boyd, D. F., Duan, S., Bajracharya, R., Awad, W. A., Apiwattanakul, N., Vogel, P., Kanneganti, T. D., and Thomas, P. G. (2018). Lung γδ T Cells Mediate Protective Responses during Neonatal Influenza Infection that Are Associated with Type 2 Immunity. Immunity 49, 531–544.e6.

Hatano, S., Tun, X., Noguchi, N., Yue, D., Yamada, H., Sun, X., Matsumoto, M., and Yoshikai, Y. (2019). Development of a new monoclonal antibody specific to mouse Vγ6 chain. Life Sci Alliance 2,

Havran, W. L., and Allison, J. P. (1990). Origin of Thy-1+ dendritic epidermal cells of adult mice from fetal thymic precursors. Nature 344, 68–70.

Havran, W. L., and Jameson, J. M. (2010). Epidermal T cells and wound healing. J Immunol 184, 5423–5428.

Herzenberg, L. A. (2000). B-1 cells: the lineage question revisited. Immunol Rev 175, 9–22.

Heyborne, K., Fu, Y. X., Kalataradi, H., Reardon, C., Roark, C., Eyster, C., Vollmer, M., Born, W., and O’Brien, R. (1993). Evidence that murine V gamma 5 and V gamma 6 gamma delta-TCR+ lymphocytes are derived from a common distinct lineage. J Immunol 151, 4523–4527.

Horton, S. J., Jaques, J., Woolthuis, C., van Dijk, J., Mesuraca, M., Huls, G., Morrone, G., Vellenga, E., and Schuringa, J. J. (2013). MLL-AF9-mediated immortalization of human hematopoietic cells along different lineages changes during ontogeny. Leukemia 27, 1116–1126.

Inlay, M. A., Serwold, T., Mosley, A., Fathman, J. W., Dimov, I. K., Seita, J., and Weissman, I. L. (2014). Identification of multipotent progenitors that emerge prior to hematopoietic stem cells in embryonic development. Stem Cell Reports 2, 457–472.

Iturri, L., Freyer, L., Biton, A., Dardenne, P., Lallemand, Y., and Gomez Perdiguero, E. (2021). Megakaryocyte production is sustained by direct differentiation from erythromyeloid progenitors in the yolk sac until midgestation. Immunity 54, 1433-1446.e5.

Kissa, K., and Herbomel, P. (2010). Blood stem cells emerge from aortic endothelium by a novel type of cell transition. Nature 464, 112–115.

Kobayashi, M., Wei, H., Yamanashi, T., Azevedo Portilho, N., Cornelius, S., Valiente, N., Nishida, C., Cheng, H., Latorre, A., Zheng, W. J., Kang, J., Seita, J., Shih, D. J., Wu, J. Q., and Yoshimoto, M. (2023). HSC-independent definitive hematopoiesis persists into adult life. Cell Rep 42, 112239.

Kondo, M., Scherer, D. C., King, A. G., Manz, M. G., and Weissman, I. L. (2001). Lymphocyte development from hematopoietic stem cells. Current Opinion in Genetics & Development Curr. Opin. Genet. Dev 11, 520–526.

Mebius, R. E., Miyamoto, T., Christensen, J., Domen, J., Cupedo, T., Weissman, I. L., and Akashi, K. (2001). The fetal liver counterpart of adult common lymphoid progenitors gives rise to all lymphoid lineages, CD45+CD4+CD3-cells, as well as macrophages. J Immunol 166, 6593–6601.

Medvinsky, A., and Dzierzak, E. (1996). Definitive hematopoiesis is autonomously initiated by the AGM region. Cell 86, 897–906.

Palis, J., Robertson, S., Kennedy, M., Wall, C., and Keller, G. (1999). Development of erythroid and myeloid progenitors in the yolk sac and embryo proper of the mouse. Development 126, 5073–5084.

Patel, S. H., Christodoulou, C., Weinreb, C., Yu, Q., da Rocha, E. L., Pepe-Mooney, B. J., Bowling, S., Li, L., Osorio, F. G., Daley, G. Q., and Camargo, F. D. (2022). Lifelong multilineage contribution by embryonic-born blood progenitors. Nature 606, 747–753.

Ramond, C., Berthault, C., Burlen-Defranoux, O., de Sousa, A. P., Guy-Grand, D., Vieira, P., Pereira, P., and Cumano, A. (2014). Two waves of distinct hematopoietic progenitor cells colonize the fetal thymus. Nat Immunol 15, 27–35.

Soares-da-Silva, F., Elsaid, R., Peixoto, M. M., Nogueira, G., Pereira, P., Bandeira, A., and Cumano, A. (2023). Assembling the layers of the hematopoietic system: A window of opportunity for thymopoiesis in the embryo. Immunol Rev 315, 54–70.

Soares-da-Silva, F., Freyer, L., Elsaid, R., Burlen-Defranoux, O., Iturri, L., Sismeiro, O., Pinto-do-Ó, P., Gomez-Perdiguero, E., and Cumano, A. (2021). Yolk sac, but not hematopoietic stem cell-derived progenitors, sustain erythropoiesis throughout murine embryonic life. J Exp Med 218,

Sörensen, I., Adams, R. H., and Gossler, A. (2009). DLL1-mediated Notch activation regulates endothelial identity in mouse fetal arteries. Blood 113, 5680–5688.

Tan, L., Fichtner, A. S., Bruni, E., Odak, I., Sandrock, I., Bubke, A., Borchers, A., Schultze-Florey, C., Koenecke, C., Förster, R., Jarek, M., von Kaisenberg, C., Schulz, A., Chu, X., Zhang, B., Li, Y., Panzer, U., Krebs, C. F., Ravens, S., and Prinz, I. (2021). A fetal wave of human type 3 effector γδ cells with restricted TCR diversity persists into adulthood. Sci Immunol 6,

Yokomizo, T., Ideue, T., Morino-Koga, S., Tham, C. Y., Sato, T., Takeda, N., Kubota, Y., Kurokawa, M., Komatsu, N., Ogawa, M., Araki, K., Osato, M., and Suda, T. (2022). Independent origins of fetal liver haematopoietic stem and progenitor cells. Nature 609, 779–784.

Yokomizo, T., and Suda, T. (2024). Development of the hematopoietic system: expanding the concept of hematopoietic stem cell-independent hematopoiesis. Trends Cell Biol 34, 161–172.

Zhang, Y., McGrath, K. E., Ayoub, E., Kingsley, P. D., Yu, H., Fegan, K., McGlynn, K. A., Rudzinskas, S., Palis, J., and Perkins, A. S. (2021). Mds1^CreERT2^, an inducible Cre allele specific to adult-repopulating hematopoietic stem cells. Cell Rep 36, 109562.

